## Supplementary Information for "Evolutionarily conserved role of serotonin signaling in regulating actomyosin contractility during morphogenesis"

$$G_{auto,i}(\tau) = \frac{\langle \delta F_i(t) \delta F_i(t+\tau) \rangle}{\langle F_i(t) \rangle^2},$$

$$G_{cross}(\tau) = \frac{\langle \delta F_g(t) \delta F_r(t+\tau) \rangle}{\langle F_g(t) \rangle \langle F_r(t) \rangle},$$

where $\delta F_i(t) = F_i(t) - \langle F_i(t) \rangle$ and $i = g, r$.

To avoid artefacts caused by long-term instabilities or single bright events, CFs were calculated segment-wise (10 segments) and then averaged. Segments showing clear distortions (typically less than 20% of all segments) were manually removed from the analysis (3).
A model for two-dimensional diffusion in the membrane and Gaussian focal volume geometry (1) was fitted to all CFs:

$$G(\tau) = \frac{1}{N}\left(1 + \frac{\tau}{\tau_d}\right)^{-1/2}\left(1 + \frac{\tau}{\tau_d S^2}\right)^{-1/2}$$

To ensure convergence of the fit for all samples (i.e. ACFs and CCFs of correlated and uncorrelated data), positive initial fit values for the particle number *N* and thus $G(\tau)$ were used. In the case of uncorrelated data, i.e. for CFs fluctuating around zero, this constraint can generate low, but positive correlation amplitudes due to noise. From the diffusion time $\tau_d$, the diffusion coefficient $D$ was determined by $D = \frac{\omega_0^2}{4\tau_d}$. The waist $\omega_0$ was determined from point FCS measurements with Alexa Fluor® 488 (Thermo Fisher Scientific, Waltham, MA, USA) dissolved in water at 20 nM, which were performed at the same laser power and 2μm depth to minimize aberrations. The structure parameter *S* was fixed to the average value determined in calibration measurements, typically around 5. From the determined particle number *N*, the protein surface concentration $c$ was quantified: $c = \frac{N}{A_{\text{eff}}} = \frac{N}{\pi\omega_0^2 S}$, where $A_{\text{eff}} = \pi\omega_0^2 S$ is the effective detection area (1).

### Extended Data Fig.1| Delayed germ-band extension in 5HT2A and 5HT2B knockout embroys

**a** GBE in 5HT2A and 5HT2B knockouts

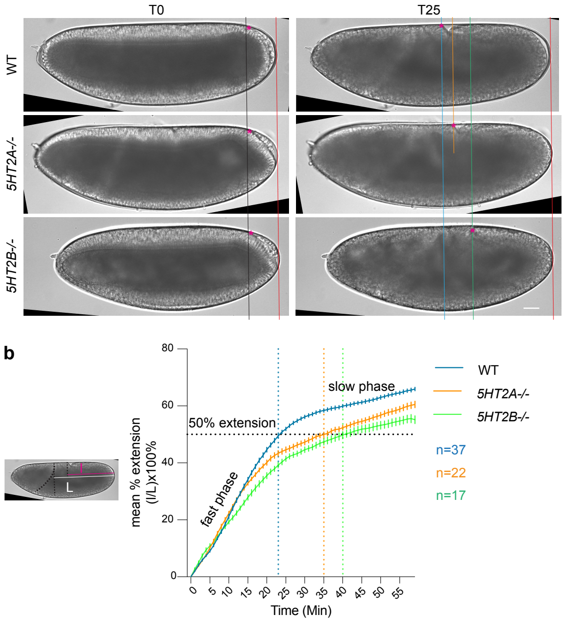

**Extended Data Fig. 1 | Delayed germ-band extension in 5HT2A and 5HT2B null mutants.** **a**) Still DIC images taken at T0 and T25 min for wild type; WT (top panels), 5HT2A null mutant; *5HT2A-/-* (middle panels) and 5HT2B null mutant; *5HT2B-/-* (bottom panels). T0 is the onset of posterior midgut rotation. The point of contact of the tissue with the vitelline membrane, marked with the pink *, is tracked. **b**) Quantification of the average distance (± s.e.m) traversed by the contact point, normalized to the maximum length it can traverse. WT embryos take ~23 minutes to achieve 50% extension, while *5HT2A-/-* and *5HT2B-/-* null mutants take ~35 and ~40 minutes respectively. Error bars SEM. Scale bar 10 µm.

**Extended Data Fig. 2 | Local tissue extension, rosettes formation, E-cad and MyoII distribution in 5HT2A loss-of-function and gain-of-funtion (Related to Figure 1).**

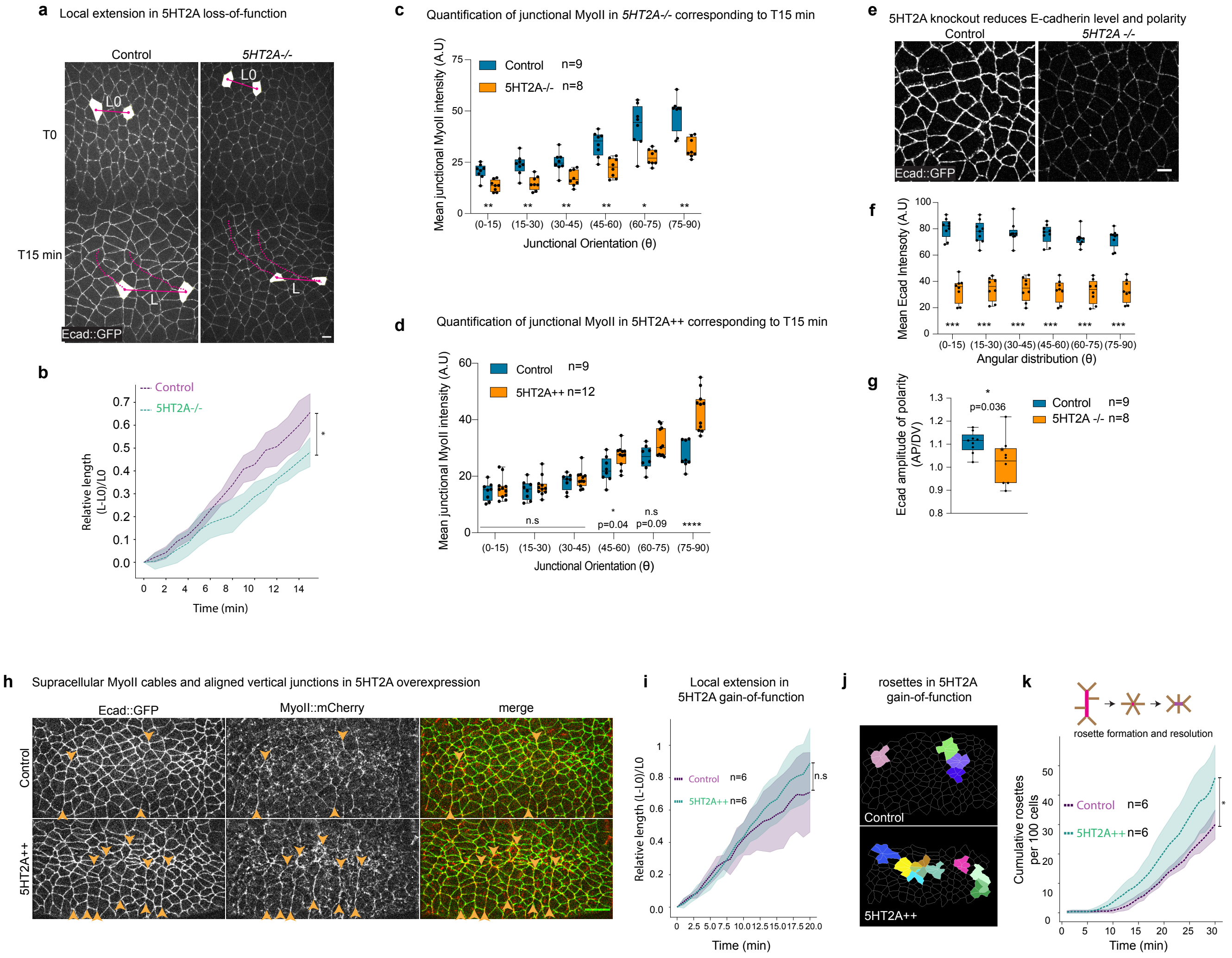

**Extended Data Fig. 2 | Local tissue extension, rosettes formation, E-cad and MyoII distribution in 5HT2A loss-of-function and gain-of-funtion (Related to Figure 1).**

**a**) Local tissue extension in control (left panels) and 5HT2A null mutant; *5HT2A-/-* (right panels) at T0 (top panels) and T15 min (bottom panels). The centroid of the cells highlighted in white are tracked for 15 minutes. The dotted pink line represents the track of the centroid of the respective white cells. **b**) Quantification of the relative length (L-L0)/L0 in different genotypes. (**c**) Quantification of MyoII intensities in different junctional orientation in control and *5HT2A-/-*. (**d**) Quantification of MyoII intensities in different junctional orientation in control and 5HT2A++. (**e-g**); **e**) Ecad::GFP in control (left panel) and *5HT2A -/-* (right panel). Quantification of Ecad::GFP levels **f**), and polarity **g**) for the respective genotypes. **h**) Snapshots of Ecad::GFP (left panel) and MyoII (middle panel) and merge of both (right panel), in control (top panels) and 5HT2A overexpression, 5HT2A++ (bottom panels). Orange arrowheads indicate the aligned cell-cell junctions forming supracellular cables. **i**) Quantification of relative length (L-L0)/L0 in the respective genotypes. **j**) Representative images of rosettes in control (top panel) and 5HT2A++ (bottom panel) taken for T30 min. **k**) Quantification of cumulative rosette counts per 100 cells in the respective conditions. ns: $p$>0.05, * $p$< 0.05, ** $p$<0.005, *** $p$<0.0005, **** $p$<0.00005, ***** $p$<0.000005 from Mann-Whitney test. n=number of embryos. Scale bars 5 µm.

**Extended Data Fig. 3 | Ectopic 5HT2A::mNeonGreen localization is not polarized yet MyoII is hyper-polarized.**

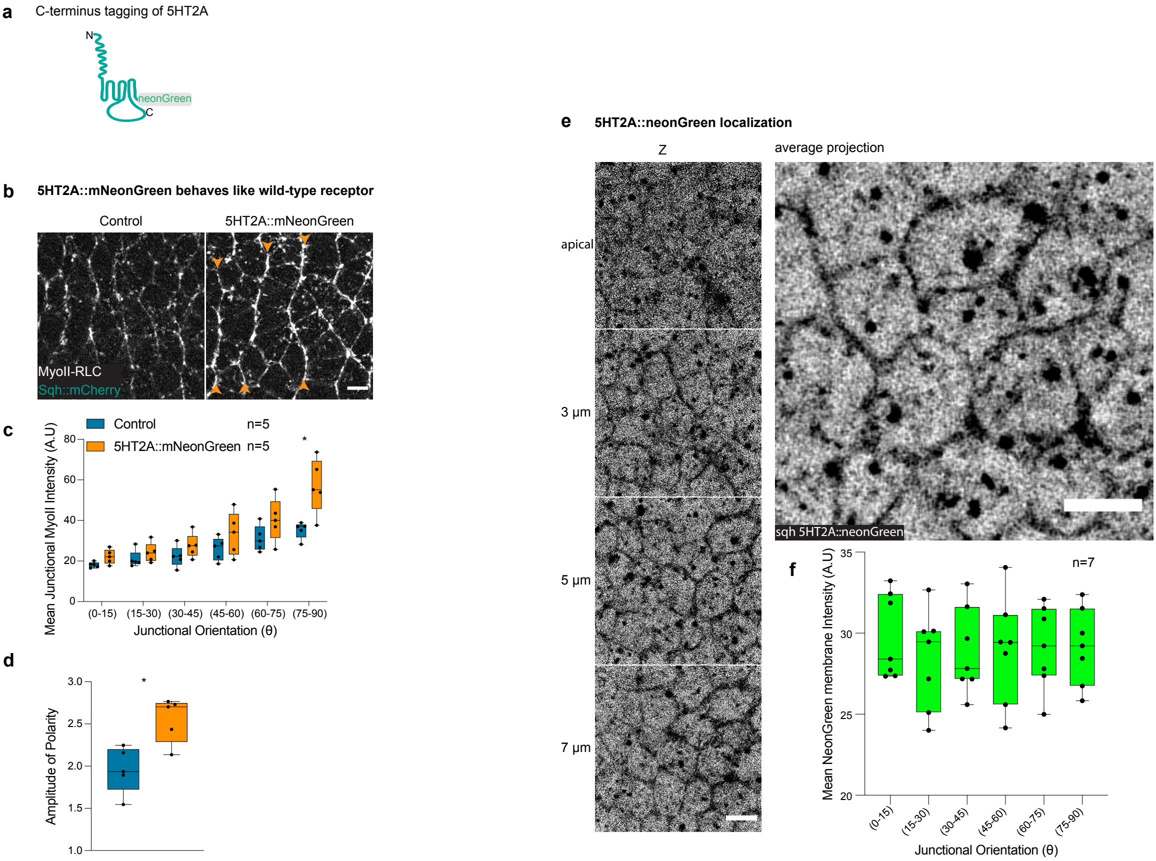

**Extended Data Fig. 3 | Ectopic 5HT2A::mNeonGreen localization is not polarized yet MyoII is hyper-polarized.**

**a**) Schematic showing C-terminal tagging of 5HT2A with mNeonGreen. (**b-d**) MyoII in 5HT2A::mNeonGreen. **b**) Still image of MyoII in control (left), 5HT2A::mNeonGreen over-expression (right, orange arrowheads indicate hyper-polarization of junctional MyoII). Quantification of junctional MyoII intensity distribution (**c**), and amplitude of polarity (**d**) in different conditions. (**e-f**) Localization of 5HT2A::mNeonGreen. Distribution of signal in different z-planes (left, from top to bottom), average projection (right) (**e**). Quantification of the distribution of membrane signal in different categories of junctions shows homogenous expression of the receptor (**f**). * $p < 0.05$ from Mann-Whitney test. n=number of embryos. Scale bars 5 μm.

**Extended Data Fig. 4 | 5HT2A::mNeonGreen and MyoII distribution following Toll-2,6,8 knockdown**

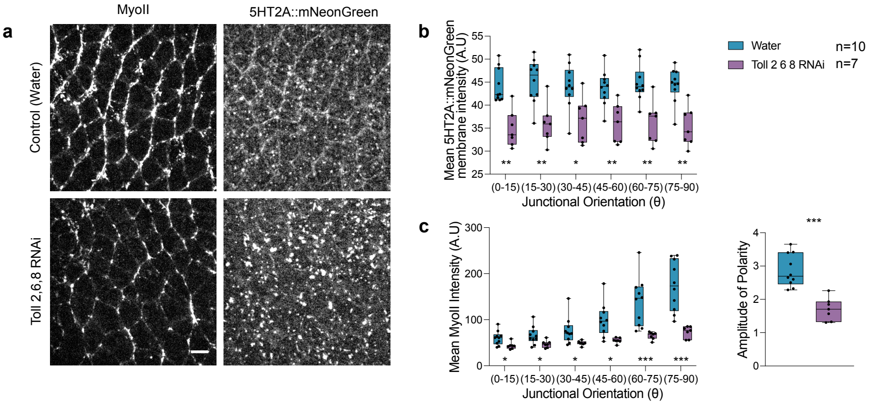

**Extended Data Fig. 4 | 5HT2A::mNeonGreen distribution following Toll-2,6,8 knockdown.**

(**a-c**) 5HT2A::mNeonGreen membrane localization following Toll-2,6,8 knock-down (Toll-2,6,8 RNAi). **a**) MyoII (top and bottom left panels) and 5HT2A::mNeonGreen (top and bottom right panels) images in water injected control (top panels) and Toll 2,6,8 RNAi injected embryos (bottom panels). Quantification of 5HT2A::mNeonGreen signal at the lateral membrane (**b**), and MyoII signal in different junction categories and amplitude of polarity (**c**) in respective conditions. * $p< 0.05$, ** $p<0.005$, *** $p<0.0005$ from Mann-Whitney test. n=number of embryos. Scale bar 5 μm.

**Extended Data Fig. 5 | Ectopic 5HT2A represses myosin phosphatase and requires Rok activity to hyper-polarize junctional MyoII.**

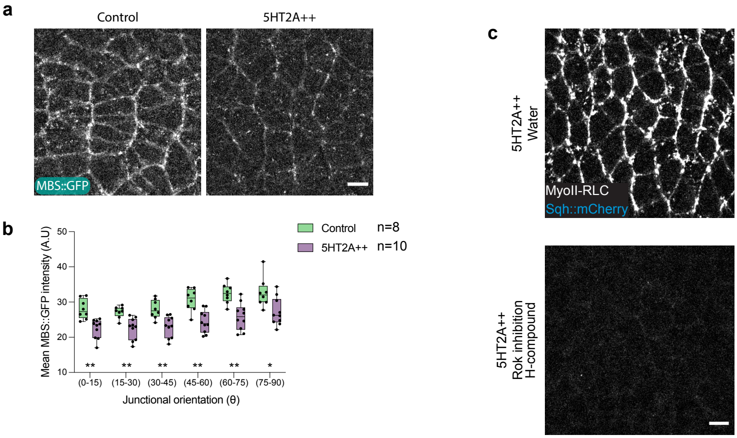

**Extended Data Fig. 5 | Ectopic 5HT2A represses myosin-phosphatase and requires Rok activity to hyper-polarize junctional MyoII.**
(**a-b**) Distribution of the myosin binding subunit (MBS::GFP) of myosin phosphatase in 5HT2A over-expression. Images of MBS::GFP in control (left panel) and 5HT2A++ (right panel) (**a**), quantification of junctional signal in respective conditions (**b**). (**c**) Rok inhibition with H-1152 compound in 5HT2A++. Snapshot of MyoII in water injected 5HT2A++ control (top panel) and H-compound injected 5HT2A++ embryo (bottom panel). ns: $p$>0.05, * $p$< 0.05, and ** $p$< 0.005 from Mann-Whitney test. n=number of embryos. Scale bars 5 μm.

**Extended Data Fig. 6 | MyoII distribution following 5HT2B knockout and knockdown.**

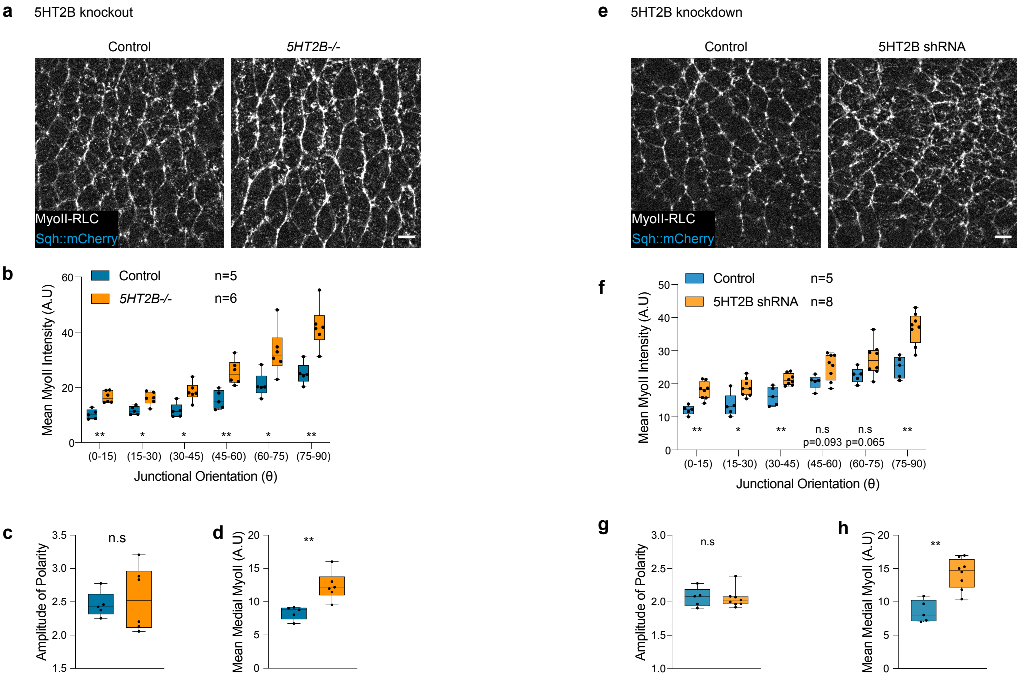

**Extended Data Fig. 6 | MyoII distribution following 5HT2B knockout and knockdown.** (**a-d**) MyoII distribution in 5HT2B null mutant embryos. **a**) Snapshots of MyoII in control (left panel) and *5HT2B-/-* null mutant (right panel) embryos. Quantification of junctional MyoII intensities (**b**), amplitude of polarity (**c**), and medial MyoII intensities (**d**) in different genotypes. (**e-h**) MyoII distribution in 5HT2B knockdown embryos. **e**) Still images of MyoII in control (left panel) and 5HT2B shRNA over-expressing embryos (right panel). Quantification of junctional MyoII intensities (**f**), amplitude of polarity (**g**), and medial MyoII intensities (**h**) in respective conditions. ns: $p>0.05$, * $p< 0.05$, and ** $p< 0.005$ from Mann-Whitney test. n=number of embryos. Scale bars 5 μm.

**Extended Data Fig. 7 | Localization of 5HT2B::mCherry and MyoII distribution.**

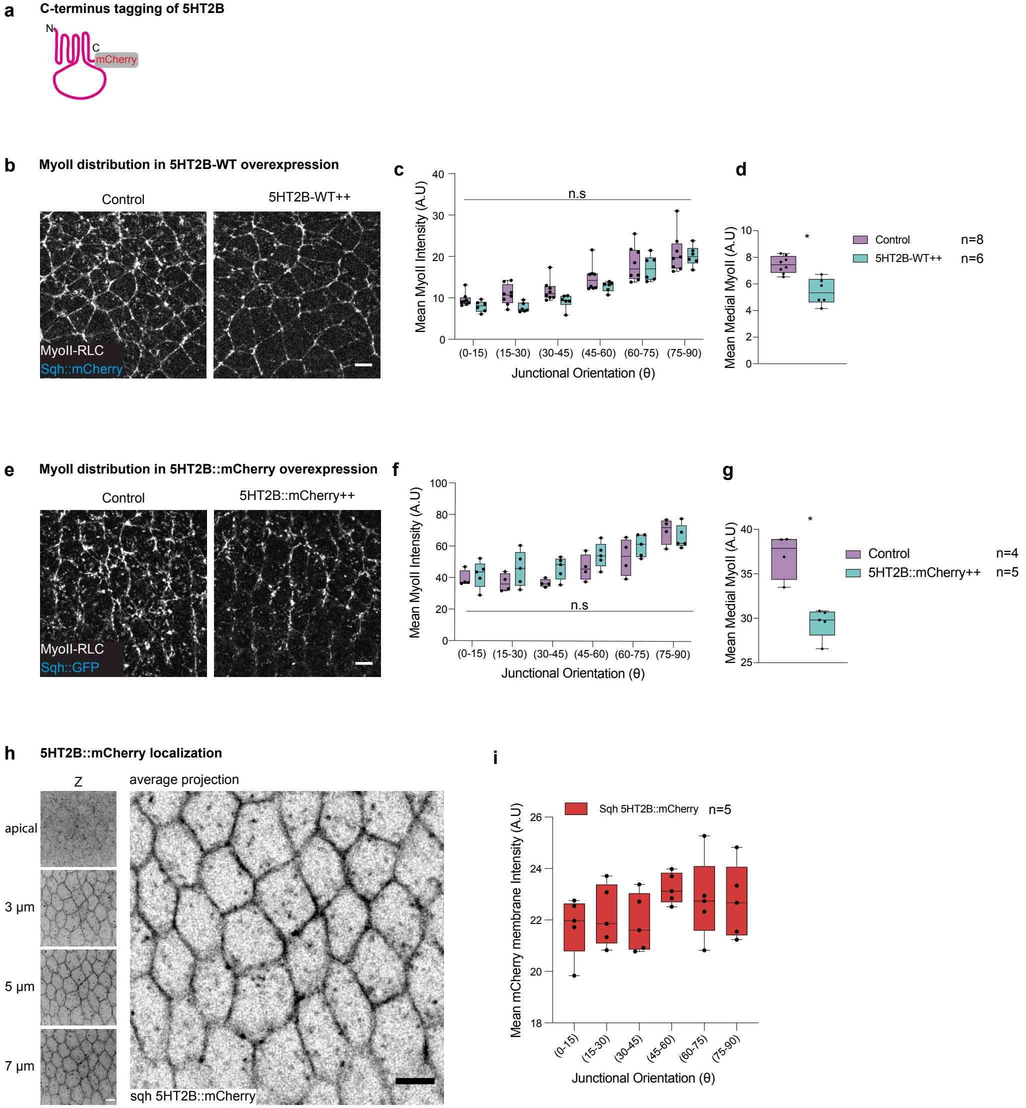

**Extended Data Fig. 7 | Localization of 5HT2B::mCherry and MyoII distribution**.
**a**) Schematic showing the C-terminus tagging of 5HT2B with mCherry. **b**) Distribution of MyoII in control (left panel) and 5HT2B-WT over-expressing (5HT2B-WT++) embryos (right panel). Quantification of MyoII at the junctions (**c**), and medial-apically (**d**) in different conditions. **e**) Images showing MyoII in control (left panel) over-expression of 5HT2B::mCherry (5HT2B::mCherry++) (right panel). Quantification of junctional (**f**), and medial MyoII intensities (**g**) in different conditions. **h**) Localization of the 5HT2B::mCherry in different z-planes (left, top-bottom apical to basal), average projection of the signal (right). **i**) Quantification of 5HT2B::mCherry membrane signal. * $p < 0.05$ from Mann-Whitney test. n=number of embryos. Scale bars 5 μm.

**Extended Data Fig. 8 | 5HT2B enhances 5HT2A endocytosis.**

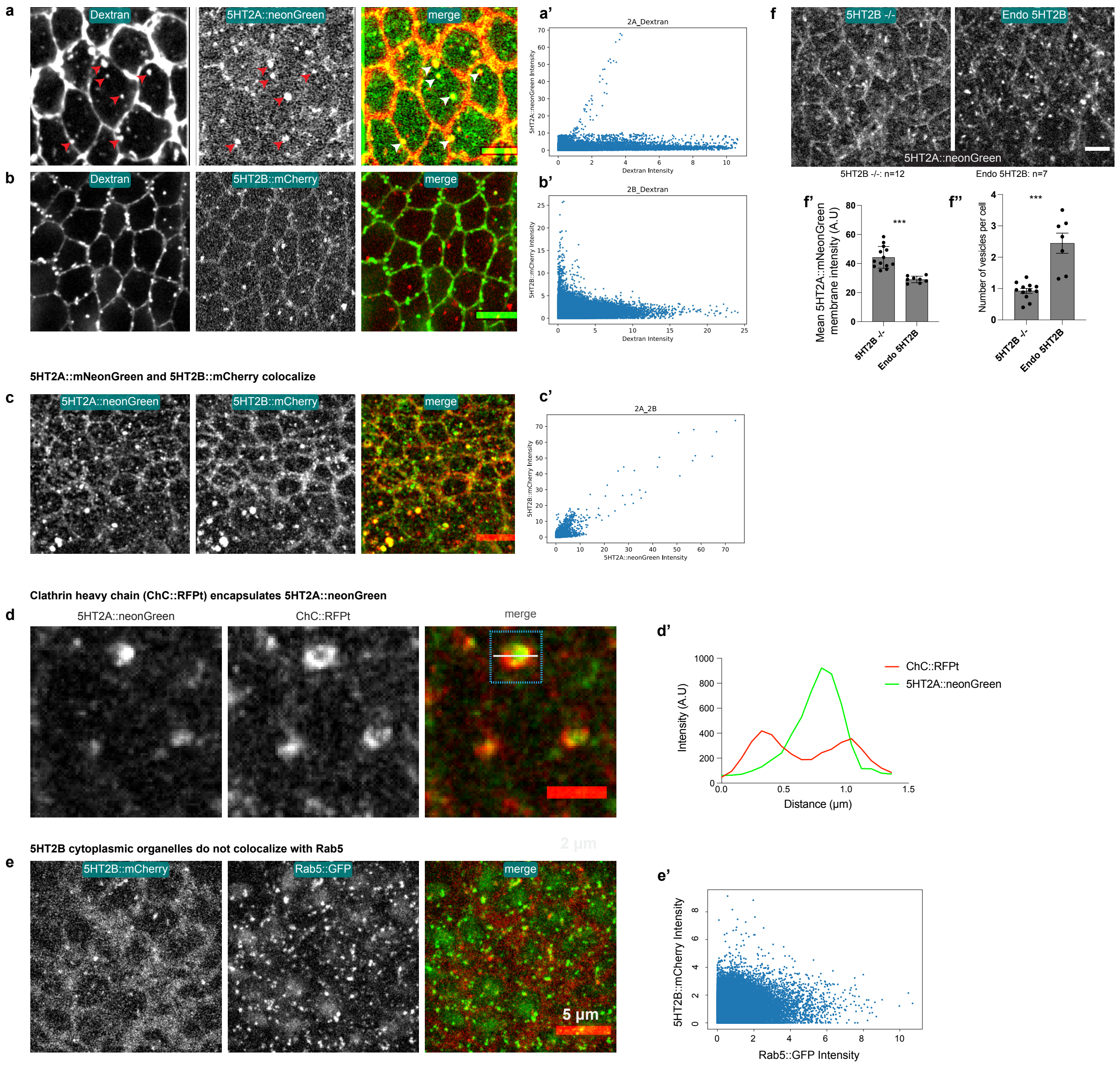

**Extended Data Fig. 8 | 5HT2B enhances 5HT2A endocytosis.**

**a**) 5HT2A::mNeonGreen co-localizes with the dextran filled mobile cytoplasmic vesicles. Dextran (left), 5HT2A::mNeonGreen (middle) and merge (right). The red and white arrowheads indicate the vesicles. (**a'**) Quantification of pixel-pixel intensity correlation between the two channels. **b**) 5HT2B::mCherry and dextran co-localization: Dextran (left), 5Ht2B::mCherry (middle) and merge (right). Fewer 5HT2B::mCherry cytoplasmic organelles co-localize with dextran filled vesicles. (**b'**) Quantification of pixel-pixel intensity correlation between the two channels. **c**) 5HT2A::mNeonGreen and 5HT2B::mCherry co-localize when co-expressed. 5HT2A::mNeonGreen (left), 5HT2B::mCherry (middle) and merge (right). (**c'**) Quantification of pixel-pixel intensity correlation between two genotypes. **d**) 5HT2A::mNeonGreen organelles encapsulated in clathrin ( ChC::RFPt) coted early endocytic organelle. 5HT2A::mNeonGreen (left), ChC::RFPt (middle) and merge (right). (**d'**) Line intensity plot across the vesicle highlighted in the blue box in the merge. **e**) 5HT2B::mCherry and Rab5::GFP marked early endosome does not co-localize. 5HT2B::mCherry (left), Rab5::GFP (middle) and merge (right). (**e'**) Quantification of pixel-pixel intensity correlation between the two channels. **f**) 5HT2A::mNeonGreen membrane levels and number of cytoplasmic organelles in 5HT2B null mutant (5HT2B-/-) embryos and WT embryos with endogenous levels of 5HT2B (Endo 5HT2B). 5HT2B-/- (left) and Endo 5HT2B (right). Quantification of the 5HT2B::mCherry membrane levels (**f'**) and the number of cytoplasmic vesicles (**f''**), in different conditions. *** $p< 0.0005$ from Mann-Whitney test. n=number of embryos. Scale bars in **a,b,c,e,f** 5 µm and in **d** 2 µm.

**Extended Data Fig. 9 | Model**

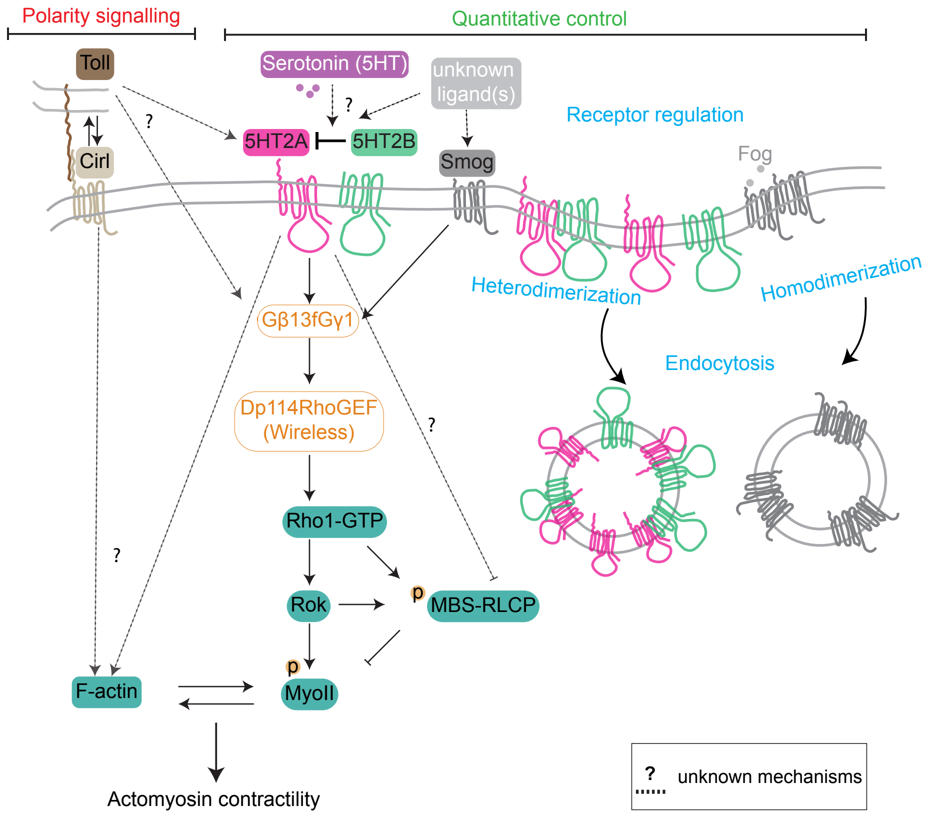

**Extended Data Fig. 9 | Model**

Model showing the modular regulation of MyoII planar polarity by Toll receptors along with the GPCR Cirl and the quantitative control of Rho1/MyoII activation by GPCRs (serotonin receptors) signalling. Serotonin/5HT2A/5HT2B regulates the junctional MyoII levels through the Gβ13f/Gγ1, Dp114RhoGEF, and Rho1 signalling pathway and signals independently of the Cirl polarity signalling. 5HT2A has a multifunctional role and regulates F-actin, activates Rho1/MyoII and inhibits myosin-phosphatase. 5HT2B inhibits 5HT2A signalling through heterodimerization and subsequent endocytosis. Toll receptors polarizes Rho1 activity via unknown GPCR(s) and F-actin through interaction with the GPCR Cirl. Cirl does not activate Rho1, but regulates F-actin and thus MyoII. Smog signalling contributes to the junctional MyoII levels. Homodimerization (Smog) and heterodimerization (5HT2A/5HT2B) and subsequent endocytosis regulate the GPCRs membrane levels and signalling. The question marks (?) and dotted arrows indicate unknown mechanisms. MBS-RLCP: Myosin Binding Subunit of Myosin Regulatory Light Chain Phosphatase. Rok: Rho-associated kinase. MyoII: MyosinII. Cirl/Latrophilin.

**Extended Data Fig. 10 | Toll receptors recruit MyoII at the junctions activating Rho1.**

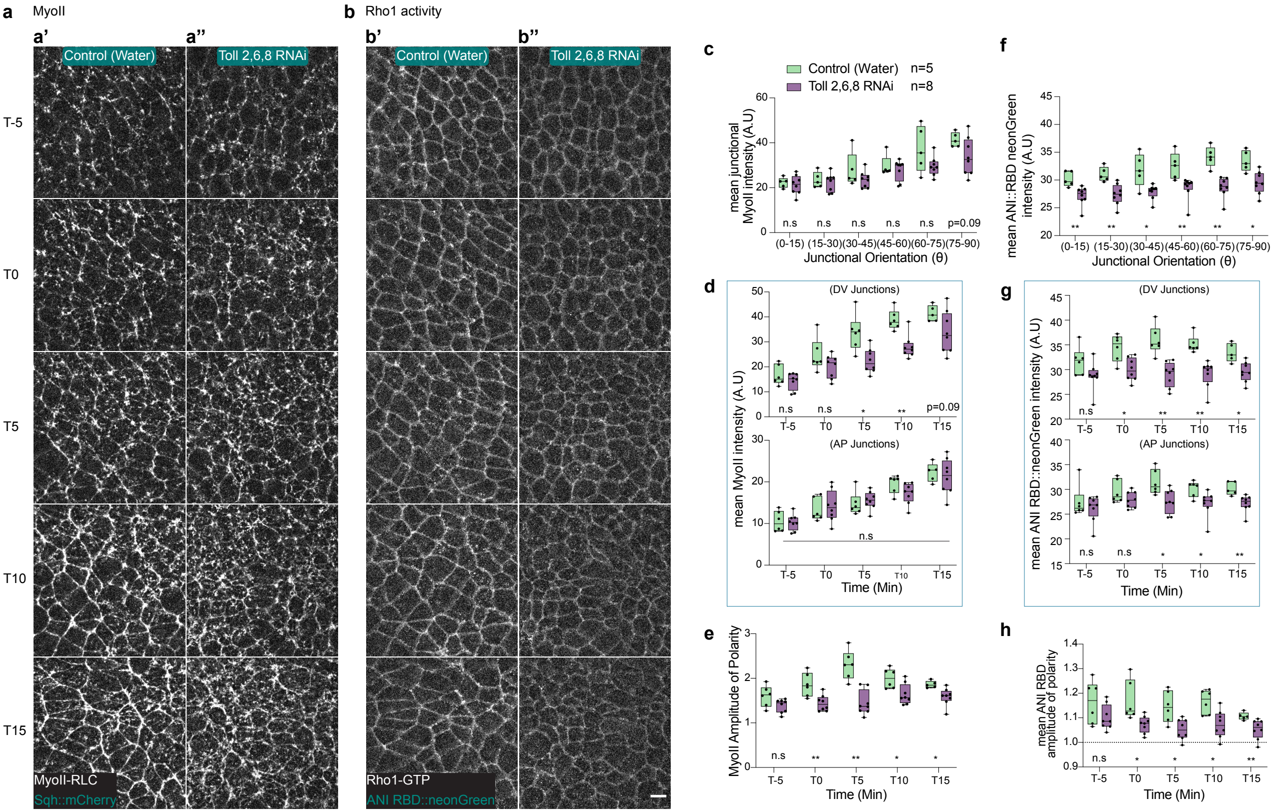

**Extended Data Fig. 10 | Toll receptors polarizes MyoII by polarizing Rho1 activity.**
(**a-h**) MyoII (**a**), and Rho1-GTP biosensor (**b**) in Toll 2,6,8 triple knock-down over-time (from top to bottom). **a**) MyoII distribution in water injected control (**a'**) and Toll-2,6,8 dsRNA (*toll*-2,6,8 RNAi) injected embryos (**a''**) over time. **b**) Rho1-GTP biosensor signal distribution in control (**b'**) and Toll-2,6,8 dsRNA (*toll*-2,6,8 RNAi) (**b''**) over time. (**c-e**) Quantification of junctional MyoII distribution taken for T15 as an example (**c**), DV and AP oriented junctions over-time (**d**), and amplitude of polarity (**e**). (**f-h**) Quantification of Rho1-GTP biosensor signal for T15 as example (**f**), DV and AP oriented junctions (**g**), and amplitude of polarity (**h**). ns: $p>0.05$, * $p< 0.05$, ** $p<0.005$ from Mann-Whitney test. n=number of embryos. Scale bar 5 μm.

**Extended Data Fig. 11 | 5HT2A and Cirl remodel F-actin.**

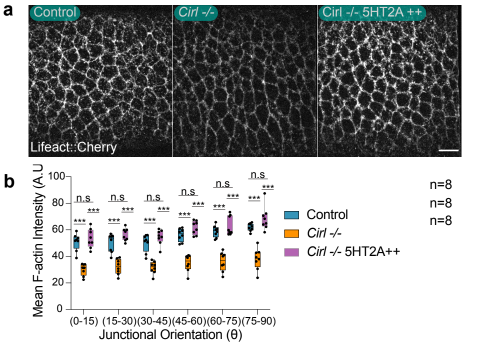

**Extended Data Fig. 11 | 5HT2A and Cirl remodel F-actin.**
**a**) Lifeact::mCherry still images in control (left panel), *Cirl -/-* (middle panel) and *Cirl-/-* and 5HT2A++ (right panel) embryos. **b**) Quantification of junctional F-actin levels in different conditions. *** $p < 0.0005$ from Mann-Whitney test. n=number of embryos. scale bar 5 μm.

**Extended Data Fig. 12 | Reduced MyoII activity after 5HT2A/2B antagonist treatment in chick embryo.**

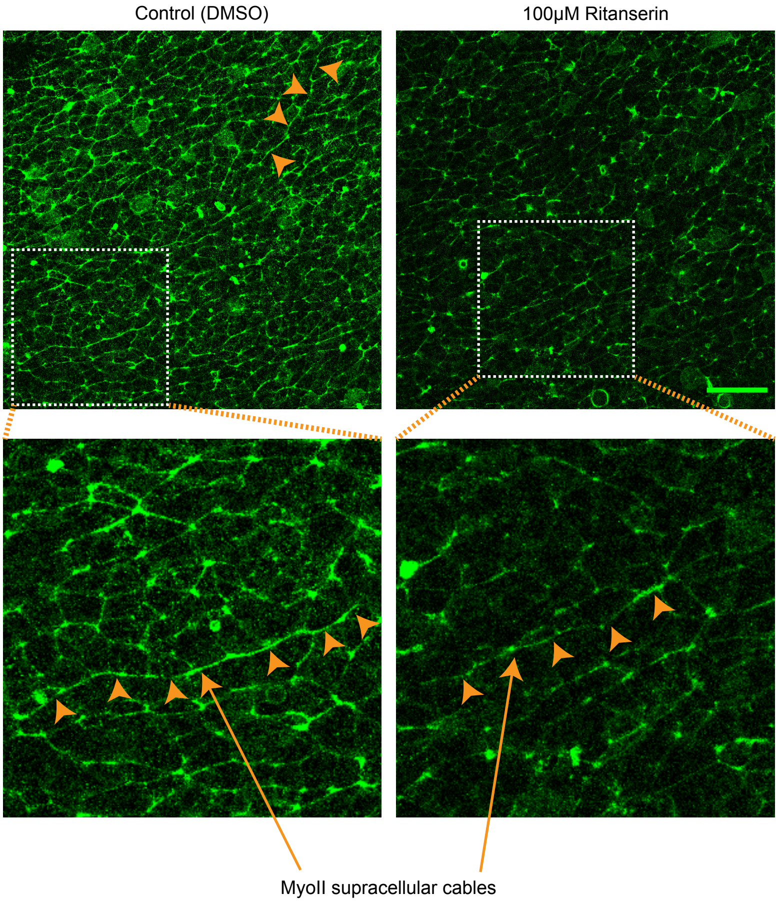

**Extended Data Fig. 12 | Reduced MyoII activity after 5HT2A/2B antagonist treatment in chick embryo.**

Images of phospho-MyoII in the embryo-proper (EP), contractile region in the DMSO treated control (left panel), 100 μM Retanserin treated (right panel) chick embryos. The white box in the images is magnified in the panels below each condition. The orange arrowheads indicate supracellular MyoII cables. The green scale bar is 50 μm.
